# A Tunable and Druggable Mechanism to Delay Forgetting of Olfactory Memories in *C. elegans*

**DOI:** 10.1101/2024.04.03.587909

**Authors:** Dana Landschaft Berliner, Kesem Goldstein, Guy Teichman, Sarit Anava, Hila Gingold, Itai Rieger, Noam Levi, Vladyslava Pechuk, Yehuda Salzberg, Priti Agarwal, Dror Sagi, Dror Cohen, Evelina Nikelshparg, Anat Ben-Zvi, Ronen Zaidel-Bar, Antonio Miranda Vizuete, Meital Oren-Suissa, Oded Rechavi

## Abstract

The poet W.B Yeats wrote that *“All that is personal soon rots, it must be packed in ice or salt”*. Here we show that in *Caenorhabditis elegans* nematodes, simple animals with just 302 neurons, memories are preserved on ice and in lithium salt. *C. elegans* nematodes can form associative memories, which are typically forgotten quickly. We discovered that when placed on ice, worms delay forgetting of specific olfactory memories by at least 8-fold. Delayed forgetting was canceled completely when the worms were gradually adapted to low temperatures, owing to a genetically-encoded program that turns acclimated worms cold-tolerant. RNA-seq, mutant analyses, and pharmacological assays revealed that regulation of membrane properties switches cold-induced delayed forgetting ON and OFF, and, remarkably, that lithium delays forgetting only in cold-sensitive but not cold-tolerant worms. We found that downregulation of the diacylglycerol pathway in the AWC sensory neurons is essential for lithium-mediated delayed forgetting, and using neuronal activity recordings located the memory trace to the downstream AIY interneurons. We suggest that the awesome genetic tractability of *C. elegans* might be harnessed to study the effects of lithium and cold temperatures on the brain, why it influences psychiatric disorders, and even more fundamentally how memory is stored and lost.

## Introduction

The nervous system switches between different internal states, and the tradeoffs between conflicting needs such as sleep-wakefulness (Gervasoni et al., 2004) or fight-flight/rest-digest (Harris & Carr, 2016, Russell & Lightman, 2019) have been studied at different levels (Briggman et al., 2005; Briggman & Kristan, 2008; Kringelbach & Deco, 2020). Imbalance between internal brain states can be pathological, for example it could cause mood disorders (Brady et al., 2017) and accelerate neurodegeneration (Saenger et al., 2017), but it can also lead to extraordinary abilities, such as Hyperthymestic syndrome (highly superior memory, Parker et al., 2006) or Savant syndrome (exceptional ability in one domain, and impairments in other domains, Treffert, 1988).

Forgetting, the flip-side of remembering, is also an adaptive process allowing organisms to flexibly respond to changing environmental conditions. While forgetting is at least in part a regulated process (Davis & Zhong, 2017), it might also occur because of tradeoffs with other functions and has costs (Lee et al., 2022; Ryan & Frankland, 2022), for instance post-traumatic stress disorder (Brewin, 2011).

The nematode *Caenorhabditis elegans* is an excellent model for studying tradeoffs in the nervous system and specifically mechanisms of behavioral plasticity (Flavell et al., 2013), as it can learn associations (Ardiel & Rankin, 2010) using a relatively simple and fully-mapped nervous system that is composed of only 302 neurons (Cook et al., 2019). The *C. elegans* genome encodes for more than a thousand olfactory receptors (Robertson & Thomas, 2006) and the worm is innately attracted to a variety of odors (e.g. butanone, benzaldehyde, pentanedione, diacetyl) (Bargmann et al., 1993). Some of the worm’s inborn attraction tendencies are reduced when odorants are paired with a short period of starvation. After such pairing, the associative memories are typically forgotten quickly so that the worms exhibit again strong attraction to the odor (Arey et al., 2018; Colbert & Bargmann, 1995; Hadziselimovic et al., 2014).

Memory performance is at least partially genetically regulated and heritable, like most other complex cognitive traits (Papassotiropoulos & de Quervain, 2011). However, the distinction between biological regulation and physical processes is vague (e.g. cellular processes that are governed by phase separation, Alberti 2017), and it remains unclear to what degree physics and environmental conditions determine memory capabilities, and in particular the kinetics of forgetting. In other words, is forgetting an active, biologically regulated process, or is memory lost passively? For example, is the rate of forgetting controlled and tuned according to the organism’s needs? Understanding how the environment affects the molecular mechanisms that mediate memory encoding and storage could have great implications, especially for alleviating pathological conditions of memory loss.

One property of the environment that might affect memory is temperature. Adapting to low temperatures requires physiological and behavioral plasticity, and indeed across phyla, different mechanisms have evolved to allow animals to sense and respond to alterations in temperatures (Angilletta & Angilletta, 2009). *C. elegans* “remembers” its cultivation temperature, and upon sensing temperature changes, modulates its temperature seeking behavior accordingly (Garrity et al., 2010; Hedgecock & Russell, 1975; Takeishi, 2022). Moreover, the worm possesses a plastic, genetically encoded mechanism that allows it to survive in very cold environments (to become cold tolerant): Adult *C. elegans* kept for more than 5 hours in 15°C can survive 48h in a temperature of 2°C, while worms which are not adapted to low temperatures and grow in 20°C or higher all die in the same timespan (Murray et al., 2007; Ohta et al., 2014; Savory et al., 2011). Ingenious studies elucidated the molecular pathways underlying this adaptation strategy and revealed that systemic cold tolerance is also in part regulated by the nervous system (Kage-Nakadai et al., 2016; Motomura et al., 2022; Okahata et al., 2016, 2019; Sonoda et al., 2016; Takagaki et al., 2020; Takeishi et al., 2020; Ujisawa et al., 2018). Further, turning the cold tolerance switch ON and OFF can have terminal consequences: Worms exposed to cold allocate vitellogenic lipids from the soma to the germline, promoting their descendant’s fitness at the expense of their own survival (Gulyas & Powell, 2022).

In this manuscript we describe a surprising observation linking forgetting and cold sensitivity which led us down a meandering path to discover two opposing brain states and a tunable mechanism for controlling memory and forgetting. While normally the worm forgets olfactory memories after 2-3 hours, we show that activating an internal switch can extend the duration of olfactory memories by at least 8-fold. We found that worms do not forget olfactory memories as long as they are kept on ice. This mechanism is canceled in cold tolerant worms. We identify the diacylglycerol (DAG) pathway as integral to cold-induced delayed forgetting, and finally, find that this internal “forgetting switch” responds to lithium, a drug used for treating bipolar disorder for decades.

## Results

### Cold delays forgetting

In an attempt to examine whether forgetting can be slowed down, we tested if cold temperatures affect memory kinetics. We conditioned worms to reduce their attraction toward the AWC-sensed odorant butanone, as previously described (see **Methods**, and Cho et al., 2016), by pairing butanone with starvation (see schemes in **Figures 1A,B**). Strikingly, while under normal conditions worms typically forget olfactory memories after 2-3 hours, we found that worms maintain the acquired memory for as long as they are kept on ice (**Figure 1C**. Similar results were obtained with two additional odorants which are also sensed by the AWC neurons, benzaldehyde and pentanedione, see **Supplementary Figure 1A,B**); Namely, the reduced odorant preference after training was maintained even after 16 hours when the worms were cooled down to ∼2°C (**Figure 1C**, even after 24 hours, however at this point the worms become very sick, **Supplementary Figure 1C**). Different lines of evidence point to the conclusion that the cool worms remain unattracted to the odor owing to the training (pairing to starvation) and not because of damage caused by the cold: (1) the naive worms (unconditioned control group) retained their normal attraction towards butanone although they were also kept on ice in the exact same way, (2) Butanone-conditioned worms that were kept on ice showed no change in preferences for benzaldehyde, a different odor which is also sensed by the AWC neurons (Bargmann et al. 1993, Colbert & Bargmann, 1995)(**Figure 1D**). In summary, we conclude that cooling worms on ice dramatically extends the retention of memory.

**Figure 1.**
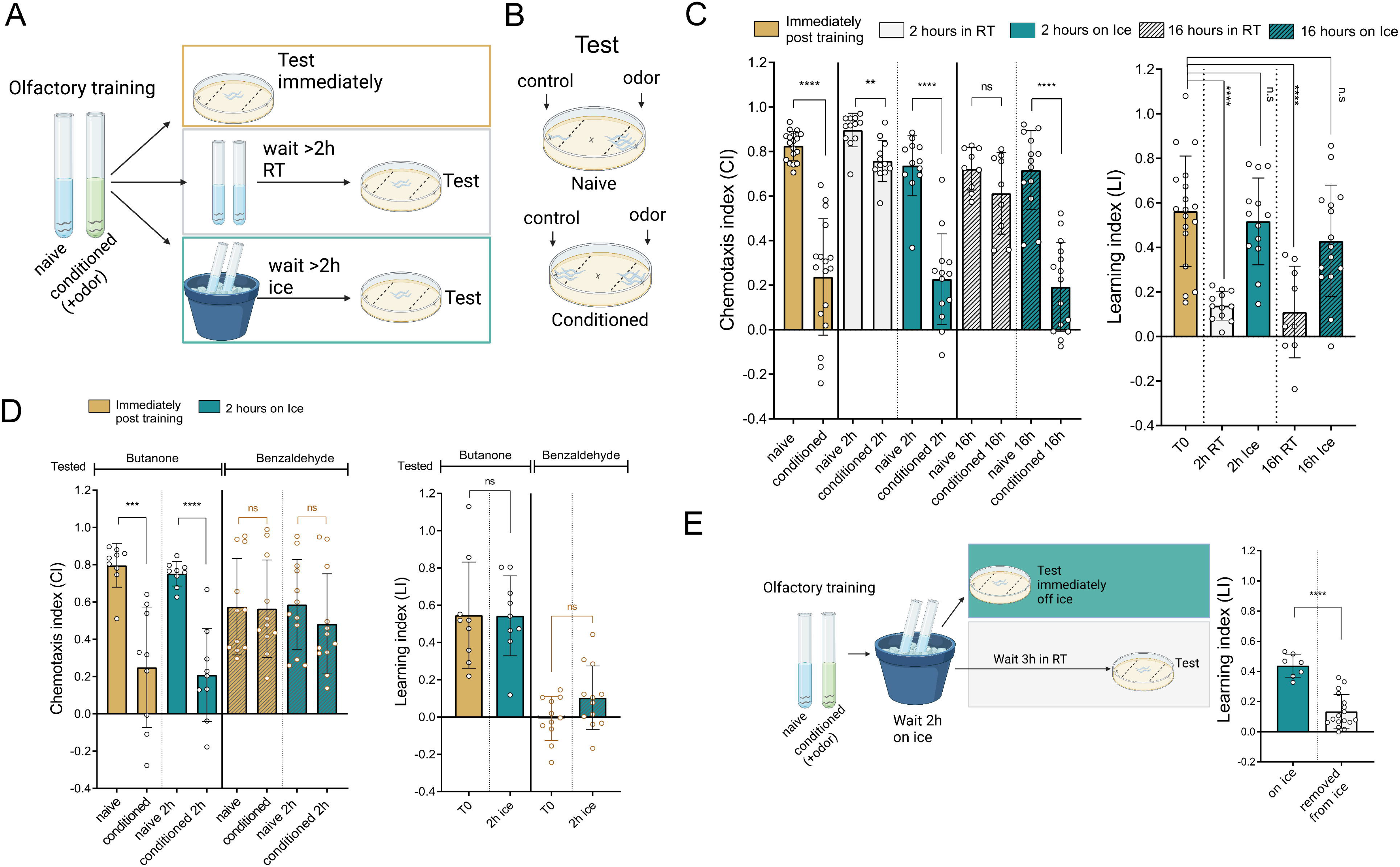
Cold Delays Forgetting. A schematic illustration of the experimental assay designed to evaluate memory retention. Adult worms were conditioned, washed and then kept either on ice or in room temperature before memory retention was tested. **B.** Graphical representation of the training outcomes, illustrating that naïve worms exhibit a stronger preference for the odor compared to conditioned (trained) worms, indicating that training succeeded. **C.** Chemotaxis outcomes for trained and control worms to the odor butanone, assessed immediately after training or after a waiting period either on ice or at room temperature (RT). The chemotaxis index (CI) and learning index (LI) serve as quantitative measures of memory retention, where CI = (#worms in odor - #worms in control area) / (total #worms in both areas), and LI = (CI of naïve group – CI of conditioned group). The left panel displays the CI, statistical significance was calculated using a two-tailed Mann-Whitney test (N from left to right: 18,17,12,12,12,13,9,9,13,15), while the right panel illustrates the LI, was calculated via one-way ANOVA followed by Tukey’s multiple comparisons test (N from left to right: 18,12,13,9,15). Each dot in the CI graph represents one chemotaxis plate (i.e. technical repeat), the graphs are comprised of 3 biological repeats with a minimum of 3 technical repeats in each. Each dot in the LI graph represents the difference between one conditioned plate and one naïve plate in the same biological repeat. **D.** Chemotaxis outcomes for trained and control worms trained with the odor butanone, assessed immediately after training or after 2 hours on ice, worms were tested for chemotaxis towards the trained odor butanone (the four leftmost bars in each panel) or for the untrained odor benzaldehyde (the four rightmost bars in each panel). In the CI (right panel) statistical significance was calculated using a two-tailed Mann-Whitney test (N from left to right: 9,9,9,9,12,12,12,12), while the LI (left panel) was calculated via one-way ANOVA followed by Tukey’s multiple comparisons test (N from left to right: 9,9,12,12). **E.** Learning index measurements for worms directly after placement on ice and 3 hours post-removal (N from left to right: 7,17) statistical significance was calculated using a two-tailed Mann-Whitney test. In panels C-E, the bar graphs denote mean ± standard deviation. Statistical significance is denoted as ns (not significant, p > 0.05), * (p < 0.05), ** (p < 0.01), *** (p < 0.001), **** (p < 0.0001).

Next, we performed experiments to distinguish between two alternative possibilities: (1) forgetting is temporarily delayed on ice (in this case the prediction is that memory will be lost quickly upon return to a normal growth temperature), or (2) cold leads to memory consolidation and long-term memory. We found that the worms forget 3 hours after they were transferred from the ice to room temperature (**Figure 1E**), and therefore conclude that cooling worms on ice delays forgetting only temporarily, and that the memory is not consolidated.

### Increasing membrane rigidity delays forgetting

Cold temperatures increase membrane rigidity (Hazel, 1995). We hypothesized that changes in membrane rigidity could lead to delayed forgetting, since the membrane’s rigidity affects neuronal activity and homeostasis (for example vesicles release dynamics, (Cossins & Prosser, 1978; Kuismanen & Saraste, 1989), and would thus extend the time it takes neurons to return to their pre-learning state.

PAQR-2 (ortholog of the adiponectin receptors AdipoR1/2) and its functional partner IGLR-2 are *C. elegans* plasma membrane proteins which detect and respond to increased membrane rigidity; Together, these proteins facilitate fatty acid desaturation, to restore membrane fluidity (Devkota et al., 2017; Svensk et al., 2013, 2016). Mutants defective in these genes were found to exhibit a highly rigid membrane.

To test the hypothesis that different factors which affect membrane rigidity, similarly to cold, may delay forgetting, we tested if the membrane rigidity sensor mutants exhibit delayed forgetting. We found that both *paqr-2* and *iglr-2* mutants delay forgetting in room temperatures (**Figure 2A,B**). These results support the hypothesis that physical hardening of the membrane slows down forgetting.

**Figure 2.**
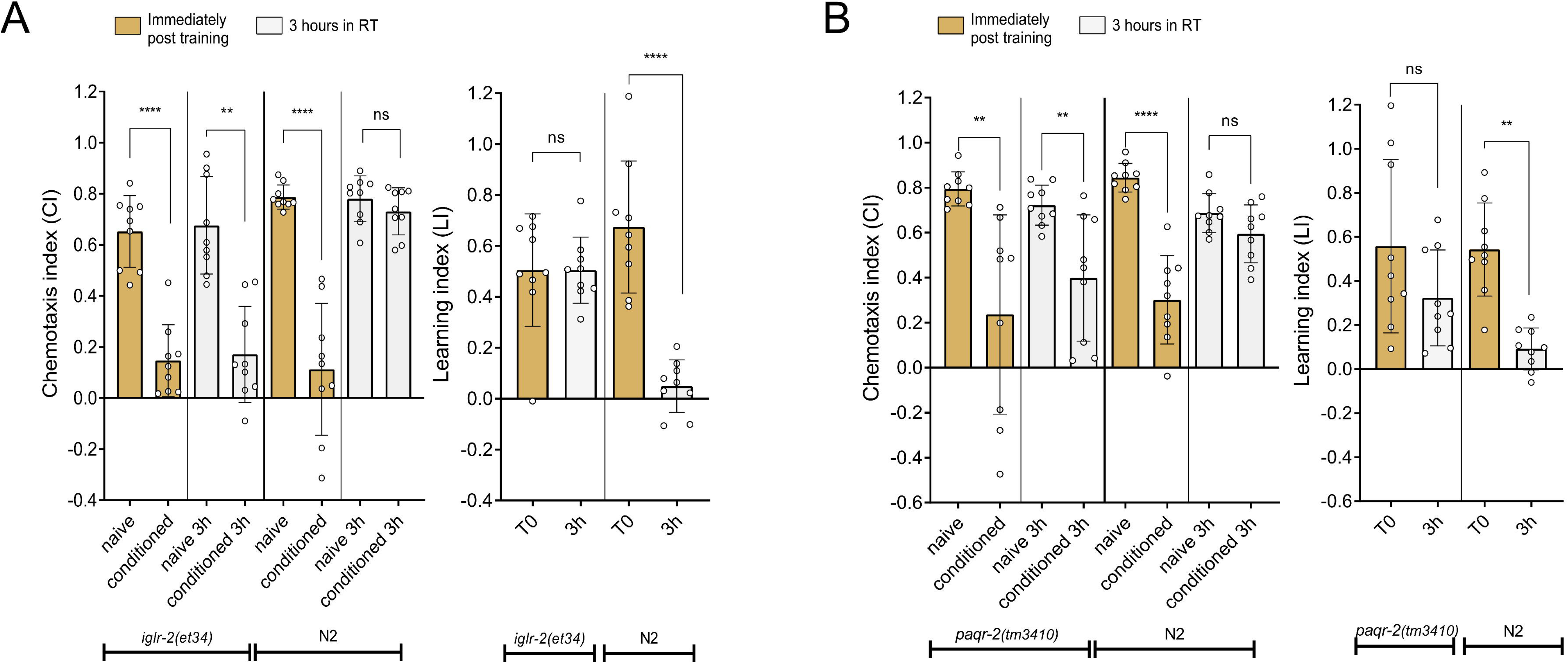
Increasing membrane rigidity delays forgetting. Chemotaxis outcomes for trained and control worms, assessed immediately after training or after a period of time at room temperature (RT) conducted on both mutant strains and N2 wild-type worms in parallel. The left panel displays the chemotaxis index (CI), statistical significance was calculated using a two-tailed Mann-Whitney test, while the right panel illustrates the learning index (LI), was calculated via one-way ANOVA followed by Tukey’s multiple comparisons test. **A.** *iglr-2*(*et34*) mutant worms. **B.** *paqr-2*(*tm3410*) mutant worms. Each dot in the CI graphs represents one chemotaxis plate (i.e. technical repeat), the graphs are comprised of 3 biological repeats with minimum of 3 technical repeat in each. Each dot in the LI graphs represents the difference between one conditioned plate and one naïve plate in the same biological repeat. Bar graphs denote mean ± standard deviation. N=9, Statistical significance is denoted as ns (not significant, p > 0.05), * (p < 0.05), ** (p < 0.01), *** (p < 0.001), **** (p < 0.0001).

### Cold tolerance cancels delayed forgetting on ice

We next examined whether the worms nevertheless possess the capacity to “resist physics” and regulate the rates of forgetting in different temperatures according to their needs. To examine this possibility, we examined forgetting rates in worms acclimated to low temperatures.

*C. elegans* become cold tolerant when acclimated to low temperatures (Murray et al., 2007; Savory et al., 2011). Specifically, Ohta et. al showed (Ohta et al., 2014) that a drastic improvement in survival in an otherwise lethal cold temperature (typically 48 h in 2°C) occurs when the cultivation temperature is changed from 20°C to 15°C (the worms must be allowed to adapt to 15°C for at least a couple of hours). Moreover, these authors discovered that the acquired cold tolerance is plastic, and is lost 2–3 hours after the cultivation temperature is changed again from 15°C to a higher cultivation temperature (Ohta et al., 2014).

We tested if there is a link between the internal processes that give rise to cold tolerance and the mechanisms that delay forgetting. We found that worms which were kept at 15°C overnight before conditioning ceased to delay forgetting on ice (**Figure 3A**). Furthermore, when worms that were cultivated overnight in 15°C were transferred back to 20°C (for 2 hours, to lose the acquired cold tolerance), they once again delayed forgetting on ice (**Figure 3B**).

**Figure 3.**
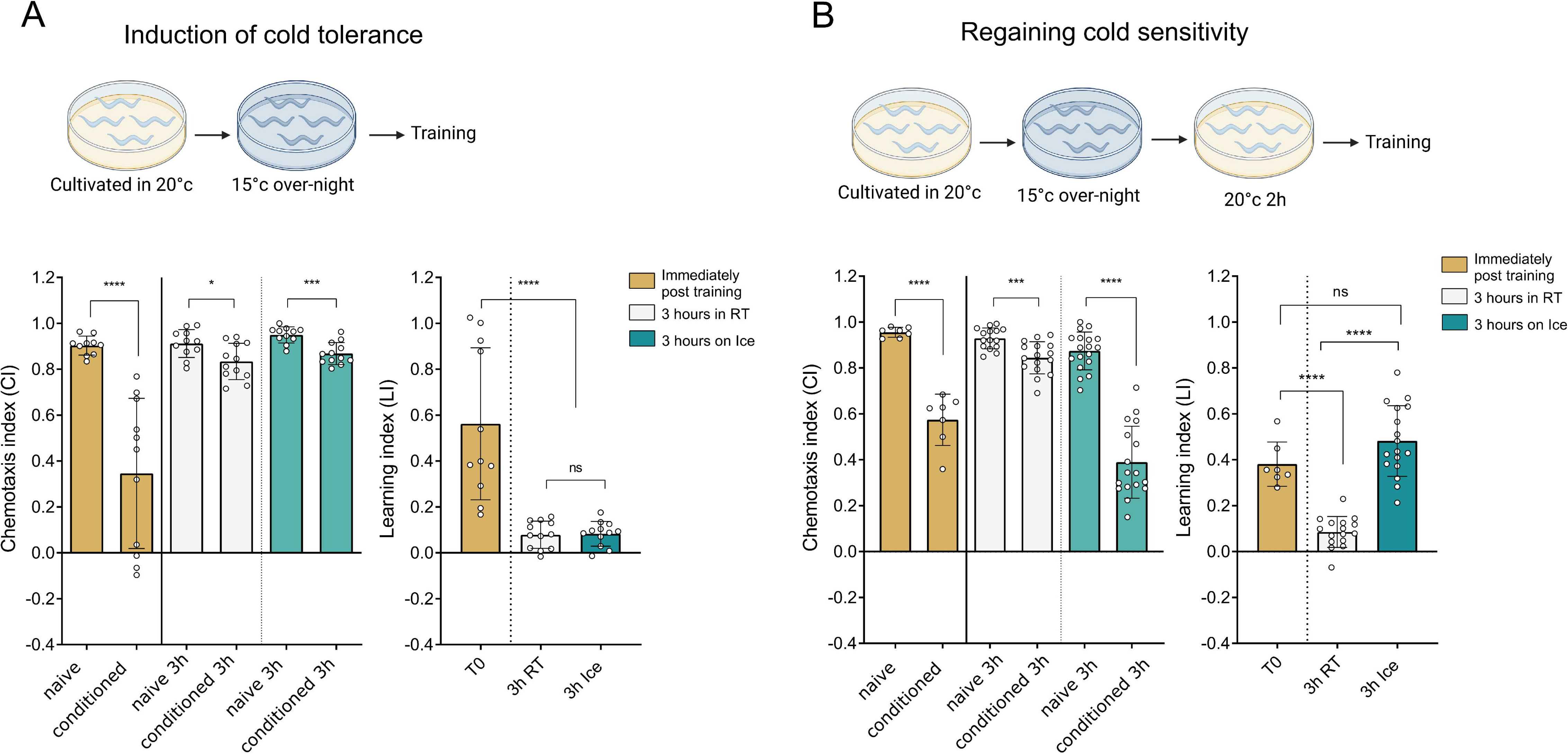
Cold tolerant worms do not delay forgetting on ice. Top: Schematic illustration of the simple procedure for induction of cold tolerance or sensitivity before memory testing. Bottom: Chemotaxis outcomes for trained and control worms, assessed immediately after training or after a period of rest either on ice or at room temperature (RT). The left panel displays the chemotaxis index (CI), statistical significance was calculated using a two-tailed Mann-Whitney test, while the right panel illustrates the learning index (LI), was calculated via one-way ANOVA followed by Tukey’s multiple comparisons test. **A**. Worms cultivated over-night in 15°C and thus rendered cold tolerant (N from left to right, CI: 10,11,11,12,11,12; LI: 11,12,12). **B.** Worm cultivated over-night in 15°C, then kept for 2 hours in 20°C to regain cold sensitivity (N from left to right, CI: 7,7,15,16,18,17; LI: 7,16,17). Each dot in the CI graph represents one chemotaxis plate (i.e technical repeat), the graphs comprised of 3 biological repeats with minimum of 3 technical repeat in each. Each dot in the LI graph represents the difference between one conditioned plate and one naïve plate in the same biological repeat. Bar graphs denote mean ± standard deviation. Statistical significance is denoted as ns (not significant, p > 0.05), * (p < 0.05), ** (p < 0.01), *** (p < 0.001), **** (p < 0.0001).

Thus, the worms can switch between two states, a cold-sensitive state during which forgetting is delayed on ice, and a cold-tolerant state, during which the memory is lost. These findings suggest that delayed forgetting on ice is a regulated mechanism, and does not occur simply because of the passive, nonspecific slowing down of molecular activity which occurs at low temperatures.

### RNA sequencing implicates the diacylglycerol pathway in delayed forgetting on ice

To identify genes that are involved in delayed forgetting on ice we leveraged the link that we identified between cold tolerance and forgetting. We sequenced mRNA from six different experimental conditions (three independent biological replications were performed): we compared the gene expression profiles of conditioned and naïve worms that were either (1) cultivated in 20°C (cold sensitive worms that delay forgetting on ice), (2) cultivated overnight in 15°C (cold tolerant worms that do not delay forgetting on ice), (3) cultivated in 15°C overnight and then transferred to 20°C for 2 hours (these worms lose their cold tolerance and become cold sensitive again, and thus delay forgetting on ice).

We identified genes which differentiate between cold-sensitive and cold-tolerant worms (See experimental design in **Figure 4A**, the bioinformatic pipeline in the **Methods**, and the complete list of genes in **Supplementary Table 1**). We hypothesized that among the 3182 up-regulated genes there might be genes whose action promotes delayed forgetting on ice, while amongst the 3284 down-regulated genes there might be genes whose action interrupts delayed forgetting on ice. While we identified genes that differentiate cold sensitive worms (that delay forgetting) and cold tolerant worms (that do not delay forgetting), no differential gene expression was observed between naïve and conditioned worms (specifically in the groups that delay forgetting on ice). This might suggest that such changes in gene expression are below the detection threshold or that memory retention depends on transcription-independent mechanisms.

**Figure 4.**
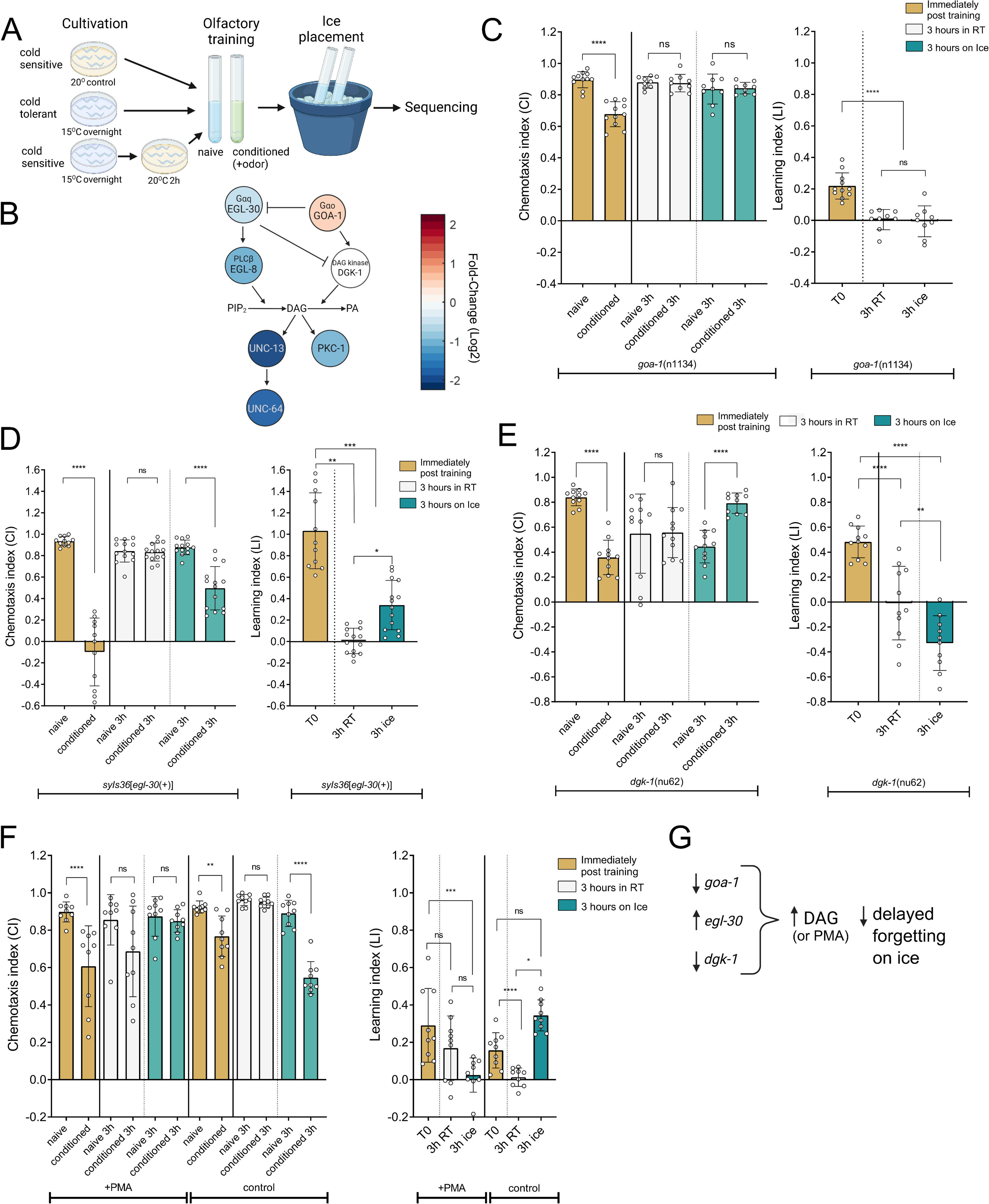
DAG antagonizes memory retention on ice. **A**. A schematic illustration of the RNA sequencing experiments. **B**. Diagram of the DAG pathway, with color code indicating the fold-change (log2 scale) in genes’ expression levels of cold-sensitive worms relative to those found in cold-tolerant worms. c-e shows chemotaxis outcomes for trained and control worms in different mutant backgrounds, assessed immediately after training or after period either on ice or at room temperature (RT). **C.** *goa-1*(*n1134*) mutants left panel: Chemotaxis Index (CI), statistical significance was calculated using a two-tailed Mann-Whitney test, right panel: Learning Index (LI), statistical significance was calculated via one-way ANOVA followed by Tukey’s multiple comparisons test (N from left to right, CI: 11,11,9,9,9,9; LI:11,9,9). **D.** *egl-30* OE strain PS2444 CI: statistical significance was calculated using a two-tailed Mann-Whitney test. LI: statistical significance was calculated via Kruskal-Wallis test followed by Dunn’s multiple comparisons test (N from left to right, CI: 11,11,13,14,13,14; LI:11,14,14). **E.** *dgk-1*(*nu62*) CI: statistical significance was calculated using two-tailed Mann-Whitney test. LI: statistical significance was calculated using one-way ANOVA followed by Tukey’s multiple comparisons test (N from left to right, CI: 11,11,11,11,11,10; LI:11,11,10). **F.** Chemotaxis outcomes of worms treated with 0.05mg/ml of the DAG analog, PMA (Phorbol-12-myristate-13-acetate) for 2 hours before training or control worms treated with ethanol as control. CI: statistical significance was calculated using two-tailed Mann-Whitney test. LI: statistical significance was calculated using one-way ANOVA followed by Tukey’s multiple comparisons test (N=9 in all groups). **G.** Diagram describing how the various mutations affect DAG levels and delayed forgetting on ice. Panels C-F: Each dot in the CI graph represents one chemotaxis plate (i.e technical repeat), the graphs comprised of 3-4 biological repeats with a minimum of 2 technical repeats in each. Each dot in the LI graph represents the difference between one conditioned plate and one naïve plate in the same biological repeat. Bar graphs denote mean ± standard deviation. Statistical significance is denoted as ns (not significant, p > 0.05), * (p < 0.05), ** (p < 0.01), *** (p < 0.001), **** (p < 0.0001).

A GO enrichment analysis revealed that the lists of enriched functions which are found for upregulated or downregulated genes in worms that delay forgetting are almost a mirror image of one another: The list of the genes which were upregulated in worms that delay forgetting on ice exhibited enrichment of genes that perform germline functions (see **Supplementary Figure 2A** and **Discussion**) and depletion of genes involved in neuronal functions. This raises the possibility that delayed forgetting results from downregulation of neuronal processes. We observed the exact opposite pattern in the list of the downregulated genes (**Supplementary Figure 2B**).

Importantly, we found that multiple genes (*goa-1, egl-30, egl-8, unc-13, pkc-1, unc-64*) in the diacylglycerol (DAG) pathway (see **figure 4B**) (Bastiani & Mendel, 2006) are differentially regulated in worms that delay forgetting on ice. DAG is known as an important regulator of different synaptic processes which are connected to memory and learning, for example vesicle priming, docking, and release (Kalyana Sundaram et al., 2023), also in *C. elegans* ( Arey et al., 2018; Matsuki et al., 2006). In particular, DAG signaling was found to be essential for adaptation to the odorants butanone, benzaldehyde, isopentanol and to salts, and also for butanone appetitive learning and mechanosensory habituation by affecting synaptic vesicles release (Arai et al., 2022; Arey et al., 2018; Kindt et al., 2007; Matsuki et al., 2006; Rahmani & Chew, 2021).

GOA-1 is encoded by an ortholog of the mammalian G_i/o_ class of Gα subunits and negatively regulates the G_αq_ protein EGL-30, whose activation induces an increase in the concentration of DAG (Miller et al., 1999). Our sequencing results indicated that *goa-1* mRNA levels are significantly up-regulated in worms that delay forgetting on ice, while *egl-30* levels are significantly down-regulated (scheme in **Figure 4B**). We thus tested forgetting in multiple mutants in this pathway, on and off ice.

We found that unlike wild type animals, *goa-1* mutants do not delay forgetting on ice (**Figure 4C**). However, it should be noted that *goa-1* animals exhibit very weak learning capabilities (as noted before by Matsuki et al., 2006). Similarly, we found that transgenic animals that over-express *egl-30* are defective in their ability to delay forgetting on ice (**Figure 4D**). Diacylglycerol kinase catalyzes the conversion of DAG to phosphatidic acid (PA) and thus mediates reduction of DAG levels in the cell membrane (Luo et al., 2004). We tested a loss-of-function allele of the diacylglycerol kinase *dgk-1* (Nurrish et al., 1999) and found that *dgk-1* mutant worms do not delay forgetting on ice. Moreover, we observed the opposite effect, as on ice trained worms showed a higher preference towards butanone, while the naive control worms exhibited a reduced attraction to the odor (**Figure 4E**).

Last, we treated worms with a DAG analog, PMA (Phorbol-12-myristate-13-acetate), to test if indeed cold delays forgetting by reducing DAG levels, and if this effect could be reversed by drugs and not only genetically. We found that supplementing the worms with the DAG analog canceled cold-induced delayed forgetting (**Figure 4F, Supplementary figure 3A**).

Together, all these experiments supported the hypothesis that DAG accumulation antagonizes cold-induced delayed forgetting (see summary in **Figure 4G**).

### Lithium delays forgetting in cold sensitive worms

The establishment of systemic cold tolerance is known to be regulated by the nervous system; More specifically, it was found that the ASJ neurons are the main regulators of this non-cell autonomous effect (Ohta et al., 2014). It was recently shown that lithium causes a selective neuronal dysfunction in the ASJ neurons (Meisel & Kim, 2016). To examine if ASJ affects forgetting, we treated the worms with lithium chloride salt (LiCl). We subjected young adult worms to 15mM of LiCl overnight, as previously described (Meisel & Kim, 2016), removed them from the lithium, and then trained them to avoid butanone. Remarkably, we found that worms which were pretreated with lithium retained the memory even five hours post training in room temperature (**Figure 5A**). Further, lithium failed to delay forgetting in cold tolerant worms (**Figure 5B**), suggesting that the mechanism by which lithium delays forgetting is linked to the mechanism by which cold delays forgetting.

**Figure 5.**
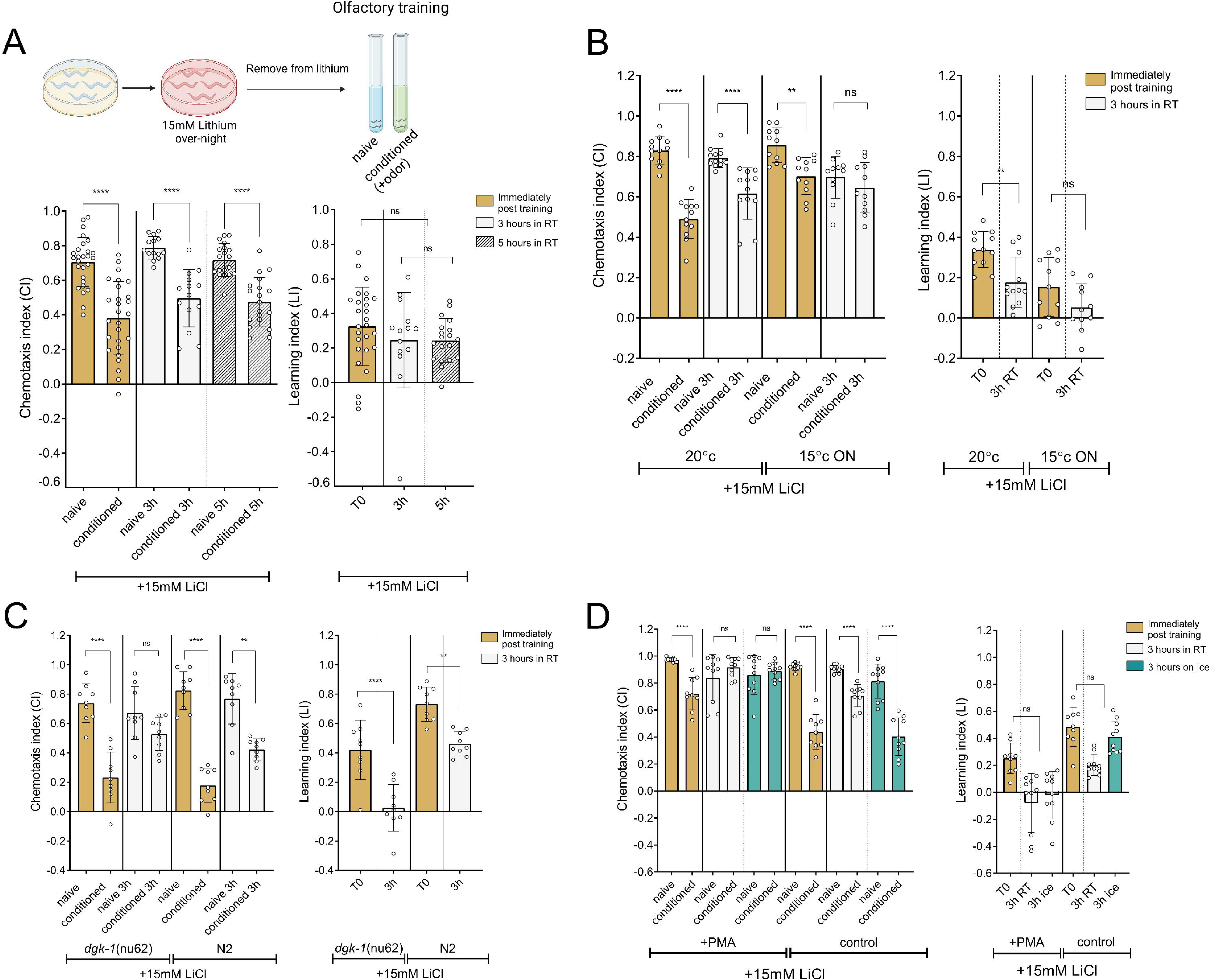
Lithium treated worms delay forgetting in room temperature. **A.** Top: schematics illustrating worms treated with lithium overnight before training. Bottom: chemotaxis outcomes for trained and control worms post lithium treatment, assessed immediately after training, after 3h or 5h at room temperature (RT). Statistical significance was calculated using two-tailed Mann-Whitney test (CI or one-way ANOVA followed by Tukey’s multiple comparisons test (LI) (N from left to right, CI:27,27, 14,14,19,19; LI:27,14,19). **B.** Chemotaxis outcomes for trained and control worms assessed immediately after training or after 3h in RT, for worms cultivated in 20°C (left) or over-night in 15°C to render them cold tolerant (right, 15°C ON) CI: Statistical significance was calculated using two-tailed Mann-Whitney test. LI: Statistical significance was calculated using one-way ANOVA followed by Tukey’s multiple comparisons test (N from left to right, CI:12,12,12,12,11,11,11,11; LI:12,12,11,11). **C.** Chemotaxis outcomes for trained and control worms in *dgk-1*(*nu62*) mutant worms (left side of each panel) or control N2 (WT), assessed immediately after training or after period either on ice or at room temperature (RT) tested in parallel CI: Statistical significance was calculated using two-tailed Mann-Whitney test. LI: Statistical significance was calculated using one-way ANOVA followed by Tukey’s multiple comparisons test (N=9). **D.** Worms treated with lithium overnight and then treated with either the DAG analog PMA or solvent (EtOH) as control during training; CI: Statistical significance was calculated using two-tailed Mann-Whitney test. LI: Statistical significance was calculated using Kruskal-Wallis followed by Dunn’s multiple comparisons test (N from left to right, CI:9,9,10,10,10,10,9,9,10,10,10,10; LI:9,10,10,9,10,10). Each dot in the CI graph represents one chemotaxis plate (i.e technical repeat), the graphs comprised of 3 biological repeats with a minimum of 3 technical repeats in each. In each panel, each dot in the LI graph represents the difference between one conditioned plate and one naïve plate in the same biological repeat. Bar graphs denote mean ± standard deviation. Statistical significance is denoted as ns (not significant, p > 0.05), * (p < 0.05), ** (p < 0.01), *** (p < 0.001), **** (p < 0.0001).

Next, we tested if lithium indeed delays forgetting by interfering with the function of the ASJ neurons, as we originally hypothesized based on lithium’s known effect over ASJ (Meisel & Kim, 2016). It was previously shown that inhibition of the ASJ neurons by lithium depends on the ASJ-specific gene *ssu-1 (Meisel & Kim, 2016)*. However, we found that lithium delays forgetting even in *ssu-1* mutants (**Supplementary Figure 3B**). Therefore, it is unlikely that lithium extends memory by inhibiting the ASJ neurons. In light of all these results, and also because we found that ASJ-ablated worms (Cornils et al., 2011) did not consistently delay forgetting (**Supplementary Figure 3C**), we reasoned that lithium delays forgetting via an ASJ-independent mechanism.

It is well established that lithium inhibits various phosphoinositol phosphates that function in inositol phosphate metabolism, such as IMPase and inositol polyphosphatase 1-phosphatase, which influence DAG turnover (Berridge et al., 1989; Brown & Tracy, 2013). Lithium’s ability to deplete inositol was suggested to be crucial for its therapeutic outcome (Harwood, 2005; Sade et al., 2016). As we showed that DAG is a key factor in the regulation of delayed forgetting, we tested whether lithium-delayed forgetting depends on DAG levels. We found that lithium-induced delayed forgetting is suppressed in *dgk*-1 mutants, in which DAG levels are elevated. (**Figure 5C**). Further, exposing lithium-treated worms to the DAG analog PMA during training prevented lithium-induced delayed forgetting (**Figure 5D**). These experiments suggest that lithium, similarly to cold shock, delays forgetting by reducing DAG. These results are intriguing also in light of lithium’s being the first line of treatment for Bipolar Disorder for more than 70 years (Cade, 1949).

### Delayed forgetting depends on reduced DAG biosynthesis in the AWC sensory neurons and is characterized by a prolonged memory trace in the AIY neurons

Next, we conducted RNA sequencing and analyzed the transcriptional changes of cold tolerant and cold sensitive worms in response to lithium (**Supplementary Table 2, Supplementary Figure 4**). We performed gene enrichment analysis and searched for genes that are differentially expressed in response to lithium treatment specifically in cold sensitive worms, as this is the condition in which worms delay forgetting.

Interestingly, our GO enrichment analysis revealed a compelling similarity in gene expression patterns between the worms that delay forgetting after lithium treatment and the worms that delay forgetting on ice. Specifically, in the group of genes that were downregulated, we observed an enrichment of genes associated with neuronal or somatic functions and depletion of germline-related terms (as observed also in cold sensitive worms on ice, see **Supplementary Figure 2**). Conversely, the group of genes that were upregulated showed an increase in germline functions (again, as did the cold sensitive worms on ice). This similarity in the transcriptional profile patterns in the worms that delay forgetting suggests a common molecular basis for these phenomena that enables memory retention.

As described above, we did not find evidence that lithium delays forgetting via its effect on the ASJ neurons (**Supplementary Figure 3A,B**). To examine if delayed forgetting is caused by a reduction in DAG levels specifically in the AWC sensory neurons, we tested *goa-1* mutants (that do not delay forgetting on ice), and *goa-1* mutants in which *goa-1*’s expression was rescued only in the AWC neurons under the control of the *odr-1* promotor (Arai et al., 2022). First, we found that rescuing *goa-1* specifically in the AWC neurons was sufficient to enable *goa-1* mutants to delay forgetting on ice (**Figure 6A**). Second, we found that while lithium treated *goa-1* mutants are completely defective in learning, rescuing *goa-1*’s expression in the AWC neurons corrects the worm’s learning ability and the capacity of the animals to delay forgetting in response to lithium (**Figure 6B**). In summary, these results suggest that maintaining low DAG levels in the AWC sensory neurons is required for delayed forgetting on ice and in response to lithium.

**Figure 6.**
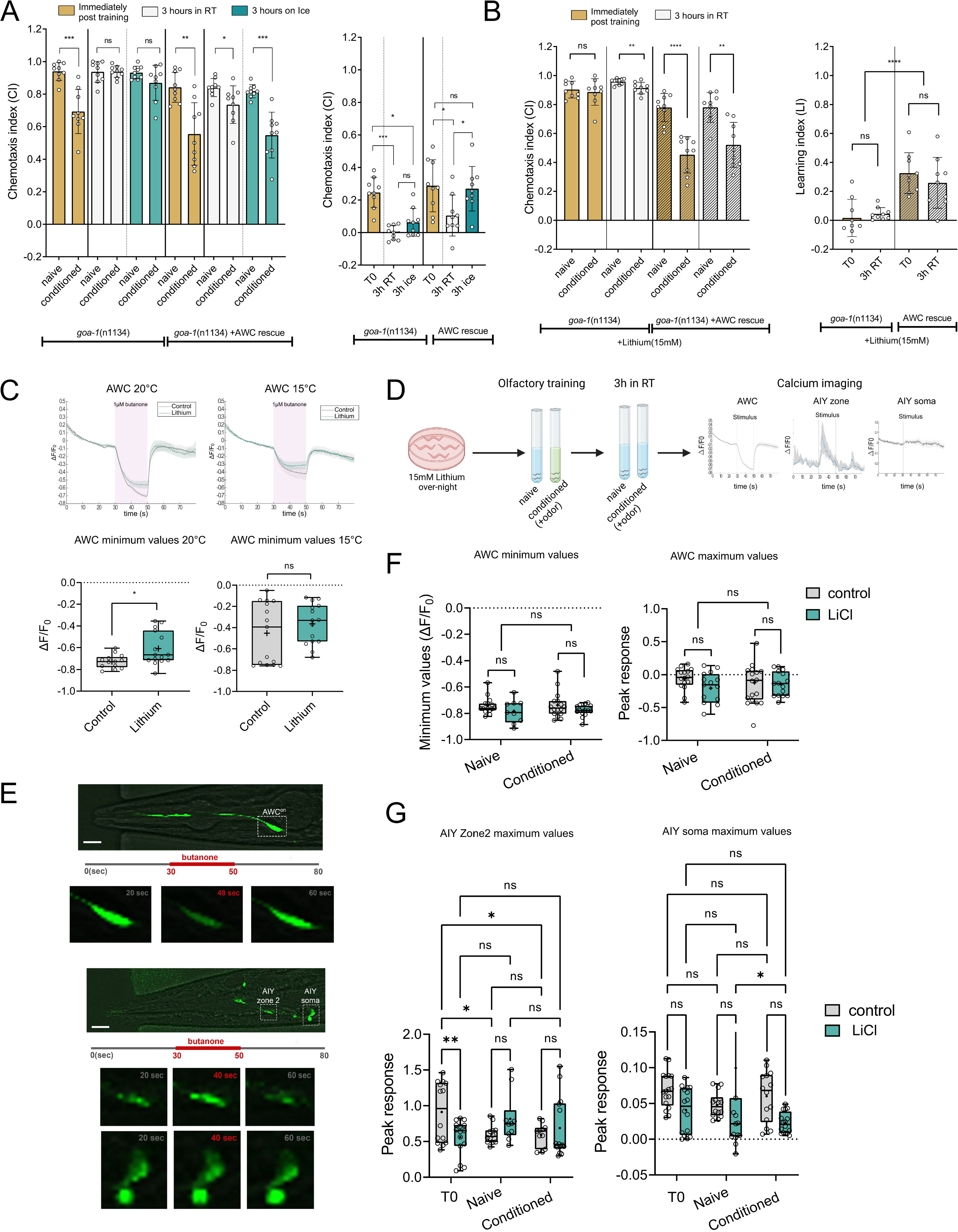
Dissecting the whereabouts of delayed forgetting. **A**. Chemotaxis outcomes for trained and control worms, assessed immediately after training or after a period of time either on ice or at room temperature (RT) conducted on both *goa-1*(*n1134*) mutant worms and in worms in which *goa-1* was rescued in AWC neurons (BFF318, *goa-1*(*n1134*);*odr-1*p::*goa*-1) in parallel. CI: Statistical significance was calculated using two-tailed Mann-Whitney test. LI: Statistical significance was calculated using one-way ANOVA followed by Tukey’s multiple comparisons test (N=9 in all groups). **B.** Chemotaxis outcomes for worms treated over-night with 15mM LiCl, trained and control worms, assessed immediately after training or after a period of time at room temperature (RT) conducted on both *goa-1*(*n1134*) mutant worms and in worms in which *goa-1* was rescued in AWC neurons (BFF318, *goa-1*(*n1134*);*odr-1*p::*goa*-1) worms in parallel. Statistical significance was calculated using two-tailed Mann-Whitney test. LI: Statistical significance was calculated using one-way ANOVA followed by Tukey’s multiple comparisons test (N=9 in all groups). Each dot in the CI graph represents one chemotaxis plate (i.e technical repeat), the graphs comprised of 3 biological repeats with a minimum of 3 technical repeat in each. Each dot in the LI graph represent the difference between one conditioned plate and one naïve plate in the same biological repeat. Bar graphs denote mean ± standard deviation. Statistical significance is denoted as ns (not significant, p > 0.05), * (p < 0.05), ** (p < 0.01), *** (p < 0.001), **** (p < 0.0001) **C.** Top: Average AWC^ON^ calcium responses to a 20-second pulse of 1µM butanone in untrained animals either treated with 15mM LiCl (turquois) or control (grey). Left panel: worms cultivated in 20°C Right panel: worms cultivated over-night in 15°C Bottom: Minimum values of the AWC neurons response shown on top panel worms cultivated either at 20°C (left, control N=13, lithium N=15) or over-night in 15 °C (right, control N=15, lithium N=15) treated with either LiCl (turquois) or control (grey), Statistical significance was calculated using two-tailed Mann-Whitney test. **D.** Schematics illustrating experimental procedures in which we tested either control or trained worms treated over-night with 15mM LiCl, for the stimulus of 1µM butanone. Three hours post training, GCaMP fluorescence was assessed at either AWC neurons or AIY neurons. **E.** Fluorescence images of GCaMP5 expressed in AWC neuron (first row) and of AWC in time lapse post exposure to the odor butanone (second row). AIY neuron (third row) GCaMP5 expressed, AIY zone and soma, post odor exposure (fourths and fifth row). **F.** Minimum (left, N from left to right: 15,16,9,15) and maximum (peak responses, right N from left to right: 8,8,15,15) values of AWC neurons in response to a stimulus of 1µM butanone, worms either treated with 15mM LiCl overnight or control, were examined 3 hours after the worms were trained to the odor butanone. **G.** Peak responses in the zone (left, N from left to right:14,15,12,10,11,14) and soma (right, N from left to right:17,16,14,15,14,16) of AIY neurons in response to a stimulus of 1µM butanone, worms either treated with 15mM LiCl overnight or control worms were examined either before training (T0) or 3 hours after the worms were trained to the odor butanone.

Next, to measure neuronal activity, we performed calcium imaging using GCaMP5s and a PDMS-based microfluidic device (Chen et al., 2013). We found that lithium reduces the response of the AWC neurons to butanone, in cold sensitive but not cold tolerant worms (**Figure 6C**). When we examined the worms 3 hours after training, in search for a “memory trace” (**Figure 6D-G**), we did not detect a significant difference in the calcium response of the AWC neurons between trained and untrained animals, or between lithium-treated and untreated worms (**Figure 6F**). Interestingly, it was observed that also another assay of olfactory training does not result in long term changes to the AWC sensory neuron activity (Chandra et al., 2023).

In contrast, and in agreement with previous studies (Chandra et al., 2023), we found that 3 hours after training, the calcium response of the AIY interneurons to butanone (sensed by the AWC neurons) is reduced in trained animals only when pre-treated with lithium. Importantly, the memory trace was evident only in the soma of the AIY neurons, but not in the axonal area known as zone 2 (**Figure 6G**), where many pre- and post-synaptic partners of AIY connect, including AWC (Colón-Ramos et al., 2007; Hawk et al., 2018).

Previous studies have shown that upon sensation of an appetitive odor, AWC inhibits the AIY interneuron, to suppress turning behavior (and thus the worms continue to move towards the odor). Aversive learning leads to a reduction in communication between the AWC and AIY neurons (Chalasani et al., 2007; Cheng et al., 2022; Tsunozaki et al., 2008; Zheng et al., 1999). Under this framework the duration of the “memory” is expected to be determined by the time it takes the AWC neurons of trained animals to resume communication with the AIY neurons. We find that the duration of this period is affected by cold sensitivity and the DAG pathway in the AWC neurons and is manifested hours after training in the AIY interneurons.

## Discussion

Our work revealed that worms have an internal, thermoregulated forgetting “switch”, which lends itself to manipulation by cold-immersion and lithium.

We found that in addition to delaying forgetting, lithium changes the way the worms move and explore their surroundings: We first noticed, by examining the tracks that the worms leave on agar plates, that even after the worms are removed from lithium, they continue to move differently. Using automated “Worm Tracking” (Javer et al., 2018, Yemini et al., 2013) we found that worms that were treated overnight with lithium display distinct behavior (**Supplementary Figure 5A,B**) and in particular elevation in angular velocity (**Supplementary Figure 5C**). Analysis of their movement, together with our memory retention assay, may be used in the future to explore lithium’s mechanism of action. In addition, we hypothesize that switching between the two “states”, which affect memory retention and movement, may also affect other traits as well. As lithium controls this transition, we speculate that the power of the *C. elegans* model might be used to study the fluctuations between internal states that occur in Bipolar Disorder (characterized also by changes in energy and activity levels, Goes, 2023).

Our study exemplifies the profound ways by which temperature affects neuronal functions (in this case, forgetting). Indeed, unsurprisingly, many studies linked cold to changes in cognitive performance (e.g. Ahlers et al., 1991; Pilcher et al., 2002; Thomas et al., 1990). It would be interesting to examine if memory loss is affected by temperature also in other animals, and in particular in humans, and more specifically whether and how lithium and the DAG pathway may influence cold-induced effects in other organisms.

Our results support an active model for forgetting and raise interesting ecological and evolutionary questions, that will hopefully be addressed in future studies. For example, why do worms forget fast, when they are perfectly capable of maintaining memories for longer periods of time? What are the tradeoff of holding memories long-term? Do the worms forget because of energy considerations or perhaps optimize some other function relevant to their ecology and life history? These are difficult questions that will hopefully keep researchers from different fields busy for many years.

## Acknowledgments

We thank all the Rechavi laboratory members for their helpful comments and fruitful discussions. We thank Oded Avishur for his roles in the project’s conceptualization and preliminary exploration. We also extend our thanks to Cornelia Bargmann, Takeshi Ishihara, Hiroshi Kagoshima, Dennis H. Kim, Joshua Meisel, Daniel Bollen and Joy Alcedo for generously sharing of worm strains. We thank Andre Brown for his advice regarding the worm’s movement following lithium treatment. We thank Rami Khosravi for helping with sequencing experiments. Some strains were provided by the CGC, which is funded by the NIH Office of Research Infrastructure Programs (P40 OD010440). D.L.B is supported partly by a fellowship from the Prajs-Drimmer Institute. O.R. is grateful for funding from the ERC (#819151), the Eric and Wendy Schmidt Fund for Strategic Innovation (Polymath Award #0140001000), the Khan Foundation (grant #0604918421), and the DFG (grant #0604918111).

## Methods

### Cultivation of worms

Worms were grown and maintained at 20°C using standard protocol (Brenner, 1974), (unless otherwise stated) on Nematode Growth Medium (NGM) plates seeded with OP50 E. coli (obtained from Caenorhabditis Genetic Center, CGC).

### Worm strains

Wild type worms were Bristol strain N2 hermaphrodites.QC136;iglr-2(et34), QC129;paqr-2(tm3410), MT2426; goa-1(n1134), PS2444; dpy-20(e1282) IV;syIs36; syIs36 [(pLB2) egl-30(+) + pBS + (pMH86) dpy-20(+)], KP1097, *dgk-1*(nu62), CX17256, *kyIs722* [*str-2*p::GCaMP5(D380Y) + *elt-2*::mCherry], Obtained from Caenorhabditis Genetic Center (CGC). CX17256*, kyIs722 [str-2p::GCaMP5(D380Y) + elt-2::mCherry]*, CX16890, kyEx4857(*mod-1*::GCaMP5; *myo-3::mCherry*], gifted by Prof. Cornelia Bargmann. BFF318; *goa-1*(n1134); qjEx30[*odr-1p*::*goa-1*+*myo-3p::gfp*] this strain was created by crossing MT2426;*goa-1*(n1134) and QD130; *goa-1(n1134)*;*tir-1*(tm3036);*qjEx30*[*odr-1p::goa-1+myo-3p::gfp*] worms, gifted by Prof. Takeshi Ishihara (QD130 used in Arai et al.,2022, see below for more details).

### Aversive olfactory learning

Gravid adult worms (10-20 individuals) were selected and placed on NGM plates seeded with OP50 to facilitate egg laying over a period of 4-6 hours, after which they were removed. Synchronized populations of worms, upon reaching day-one of adulthood, were harvested using S-basal buffer (0.1M NaCl, 0.05M KPO4) in preparation for subsequent training procedure.

### Butanone training

The protocol was adapted with modifications from Cho et al. (2016). Briefly, day-one adult worms were rinsed three times with S-basal buffer and transferred to 15 ml glass vials (20×150mm, Glassco), each containing 3ml of S-basal buffer. For the conditioning process, 30 µl of butanone diluted to 100mM in ethanol (resulting in 1mM butanone in training vial) was added to the buffer for the conditioned group, while the naive group received 30µl of ethanol. Subsequently, the worms were maintained in the liquid for either 90 minutes or, specifically for the cold tolerance assays, 45 minutes. This adjustment in conditioning time to 45 minutes for the cold tolerance experiments was necessary because cold tolerance diminishes after 2 hours at room temperature. Post-conditioning, worms were washed three times with S-basal buffer and allocated into designated groups for further experimental assays, including ice placement or room temperature maintenance.

### Room temperature maintenance

Following conditioning, worms designated for room temperature conditions were transferred to 15 ml glass vials (20×150 mm, Glassco) containing 2 ml of S-basal buffer. These vials were maintained at ambient laboratory conditions. Control conditions for extended exposure (16 hours and 24 hours): The control group for the 16-hour duration was incubated at a stable 20°C, utilizing a temperature-controlled incubator to ensure environmental consistency.

### Ice placement

For the cohorts assigned to cold exposure, 15 ml glass vials (20×150 mm, Glassco) prefilled with 2 ml of S-basal buffer were pre-cooled on ice prior to worm transfer. Worms, after undergoing three washes with S-basal buffer, were then placed into the chilled vials. For short-term ice exposure, vials were kept directly on ice. For extended cold treatment (of 16 or 24 hours) the vials were positioned within a large cooler filled with ice, which was subsequently stored in a refrigerator set to 4°C.

### Chemotaxis to tested odors

was evaluated using 90 mm diameter plates filled with 14 mL of chemotaxis agar (comprising 2% agar, 5 mM phosphate buffer at pH 6.0, 1 mM CaCl2, and 1 mM MgSO4), prepared 24 hours prior to the assay. Prior to initiating chemotaxis, worms were washed once with a chemotaxis buffer (5 mM phosphate buffer pH 6.0, 1 mM CaCl2, 1 mM MgSO4). For the assay, two 1 µl aliquots of the odorant (diluted 1:1000 in ethanol) and an ethanol control were spotted on opposite sides of the plate. Additionally, 1 µl of 1 M sodium azide (NaN3) was placed adjacent to the odorant and control spots to immobilize animals upon reaching the target area, ensuring accurate counting. Worms were centrally placed on the chemotaxis plate, and excess liquid was gently removed using Kimwipes. The assay commenced with populations ranging from 50 to 400 individuals per plate. Each experiment was conducted at least three times. Plates with fewer than 50 animals reaching the designated odor locations at the assay’s conclusion were omitted from the final analysis. Quantification involved counting the number of worms at the odorant versus control spots, with the chemotaxis index (CI) calculated as [#worms in Odor - #worms in Control] / [#worms in Odor + #worms in Control]. The Learning Index (LI), indicating the differential response between naive and conditioned populations within the same technical replicate, was determined by LI = CI_naive_ – CI _conditioned_.

### Quantification and analysis of behavioural differences

Behavioral assay data were analyzed focusing on two primary metrics: the Chemotaxis Index (CI) and the Learning Index (LI). CI quantifies the preference of the worm population towards the odor presented on each assay plate. To assess the significance of CI differences, conditioned groups were compared to their corresponding naive groups at each experimental time point. Statistical analysis was chosen based on data distribution and model assumptions: a t-test was applied for normally distributed data and variance equality, while Mann-Whitney test was used for data not meeting these criteria. LI represents the behavioral change attributed to conditioning, calculated as the difference in CI between conditioned and naive groups at each time point. For evaluations encompassing three or more time points where data adhered to statistical prerequisites, one-way ANOVA with subsequent Tukey’s multiple comparison test was conducted. In cases where data did not fit one-way ANOVA conditions, the Kruskal-Wallis test, followed by Dunn’s multiple comparisons test, was applied. For binary time point analyses, the choice between t-test and the Mann-Whitney test was dictated by the data’s distribution characteristics. All statistical analyses and graphing were performed using GraphPad Prism software.

### Graphics

Figures and schemes were designed with the aid of BioRender.com; figure alignment was done using Inkscape vector graphics editor.

### Lithium treatment

Young adult worms were harvested from standard NGM plates and subjected to an overnight (∼16 hours) treatment by placing them on NGM plates supplemented with 15 mM Lithium Chloride (LiCl, Catalogue No. L9650, Sigma-Aldrich), lithium treatment protocol was adapted from Meisel & Kim et al., 2016,. These plates were pre-seeded with *Escherichia coli* OP50 bacteria 1-2 days prior to the initiation of lithium treatment, ensuring optimal conditions for the assay.

### PMA administration

Synchronized adult worms were exposed to Phorbol 12-myristate 13-acetate (PMA, 0.05 mg/ml, Catalog No. P8139, Sigma-Aldrich) or an ethanol solvent as a control. This exposure occurred under two distinct conditions: either for a 2-hour period prior to the conditioning step when placed on NGM plates infused with PMA, or concurrently during the 90-minute conditioning phase placed inside S-basal buffer in 15 ml glass vials (20×150 mm, Glassco). During conditioning, worms were presented with either 1 mM butanone (for the conditioned group) or an equivalent volume of ethanol (for the naive group) in S-basal buffer, as outlined in the section detailing aversive olfactory training.

### *goa-1* rescue in AWC

The following strain was created by crossing *goa-1*(n1134, MT2426) to QD130 (*goa-1(n1134);tir-1(tm3036)*;*qjEx30[odr-1p::goa-1+myo-3p::gfp*]]) kindly gifted by Takeshi Ishihara lab (Arai et al., 2022). To identify and isolate *goa-1* The following primers were used followed by Sanger sequencing 5’-AGCTGCACCACATACAGTGA-3’, 5’-TCGGACGTTCTATGGGACAA-3’. To identify *tir-1*(tm3036) the following primers were used: 5’-AAGTGGCCTCCTCTCCAGAC-3’, 5’-GCACCCAATCCTCACAGTTA-3’.

### mRNA library preparation and sequencing, Exploring the transcriptional basis of delayed forgetting on ice

Worms were first trained with the odorant butanone, then place on ice for 2-hours before RNA extraction using Trizol (worms were placed in Trizol immediately upon removal from ice). mRNA sequencing was performed to investigate gene expression profiles under six distinct experimental conditions as described in **Figure 4a**. The worms were either trained or not, and the 3 different cultivation regimes were as follows: (1) Worms cultivated at 20°C, representing the group of cold-sensitive individuals that exhibit delayed forgetting when exposed to ice (2) Worms cultivated overnight at 15°C, cold-tolerant individuals that do not delay forgetting on ice. (3) Worms initially cultivated at 15°C overnight before being transferred to 20°C for 2 hours, a treatment intended to reverse the cold tolerance acquired from the overnight exposure.

cDNA was prepared with either the SMART-Seq HT Kit or the SMART-Seq v4 Ultra. A total of 10 ng RNA was used as input and processed according to the manufacturer’s protocol. Double strand cDNA was amplified by a 10 cycle PCR for SMART-Seq HT samples and 12 cycles for SMART-Seq v4 Ultra samples. cDNA quantity and quality were verified using a TapeStation 2200 (Agilent). One ng of cDNA was used as input for preparation of sequencing libraries using the Nextera XT DNA Sequencing kit (Illumina). Libraries were prepared according to the manufacturer’s instructions. Quality and concentration were verified with TapeStation 2200 and libraries were pooled according to molarity for sequencing on the NextSeq 500.

### Exploring the transcriptional basis of delayed forgetting post lithium treatment

Worms were placed overnight on plates containing either 15mM LiCl for the experimental group or NGM as control. The worms have been cultivated overnight at one of two temperatures: 15°C, to induce cold tolerance, or 20°C, to keep the worms cold sensitive. Worms were then collected in Trizol for RNA extraction. RNA libraries were prepared using the NEBNext Ultra II directional RNA library prep kit for illumina according to the manufacturer’s protocol. Quality and concentration were verified with TapeStation 2200 and libraries were pooled according to molarity for sequencing on the NextSeq 500.

### mRNA sequencing data analysis

The output FASTQ files were assessed for quality with FastQC (Andrews and Others 2010). Reads were aligned to version *ce11* of the *C. elegans* genome using HISAT2 version 2.0.5 (Sirén et al. 2014). The aligned reads were next counted and assigned to genes using the python-based script HTSeq-count (Anders et al., 2015) and the Ensembl-provided gff file (release 105), using the “intersection-nonempty” mode and ignoring secondary alignments. Next, we used the HTSeq output as an input file for differential expression with R Deseq2 (Love et al. 2014). Genes were regarded as differentially expressed if they passed the criterion ofFDR <= 0.1. Further analysis was performed using the python-based package RNAlysis (Teichman et al. 2023, Python package version 2.1.1).

### Calcium imaging and analysis

For AWC imaging: CX17256 worms were employed to investigate the response of AWC neurons to butanone exposure, following lithium treatment. Microfluidic chip fabrication and calcium imaging setup was performed as described (Pechuk et al., 2022). Worms were recorded in response to a stimulus of 1µM butanone. A minimum of 15 animals were tested in each group. Imaging was carried out using a Zeiss LSM 880 confocal microscope equipped with a 40x magnification water objective. Imaging frequency was 6.6667 Hz, total imaging duration was 80 sec, and stimulus duration was 20 sec. The fluorescence intensity of GCaMP5a was measured using FIJI. Regions Of Interest (ROIs) for the cell bodies were manually drawn and the mean of the grey values were measured. MATLAB was used for downstream data processing. For each worm individually the baseline fluorescent level (F_0_) was calculated by averaging the mean grey value of 185 frames before stimulus delivery. For each frame, ΔF was obtained by subtracting F_0_ from the value of that frame. This value was then divided by F_0_, thereby normalizing individual fluorescence baseline levels and fluctuation variability (ΔF/ F_0_). For AWC imaging post training: CX17256 worms that were cultivated overnight on either 15mM lithium plates or control plates (see lithium treatment section) were trained for the odor butanone (see aversive olfactory training section), then kept at RT for 2 hours before assessment. For AIY imaging post training: CX16890 worms were cultivated overnight on either 15mM lithium plates or control plates (see lithium treatment section), a fraction of the population was tested as baseline, while another fraction was trained for the odor butanone (see aversive olfactory training section), then kept at RT for 2 hours before assessment.

## Data and materials availability

All data needed to evaluate the conclusions in the paper are present in the paper and/or the Supplementary Materials and will be made available.

