## Supplementary figures and images for "A Tunable and Druggable Mechanism to Delay Forgetting of Olfactory Memories in *C. elegans*"

### Supplementary Figure1

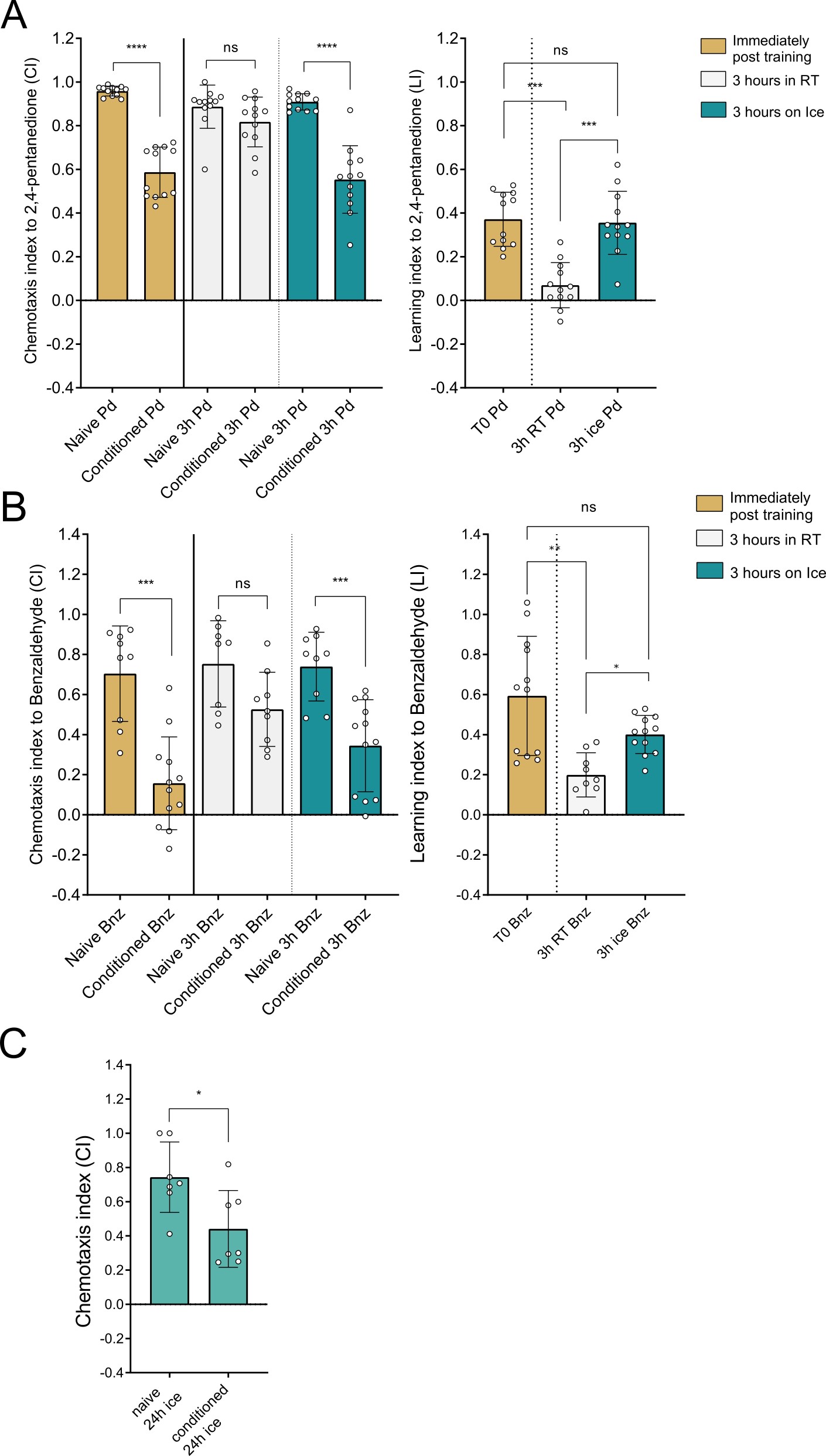

### Supplementary Figure2

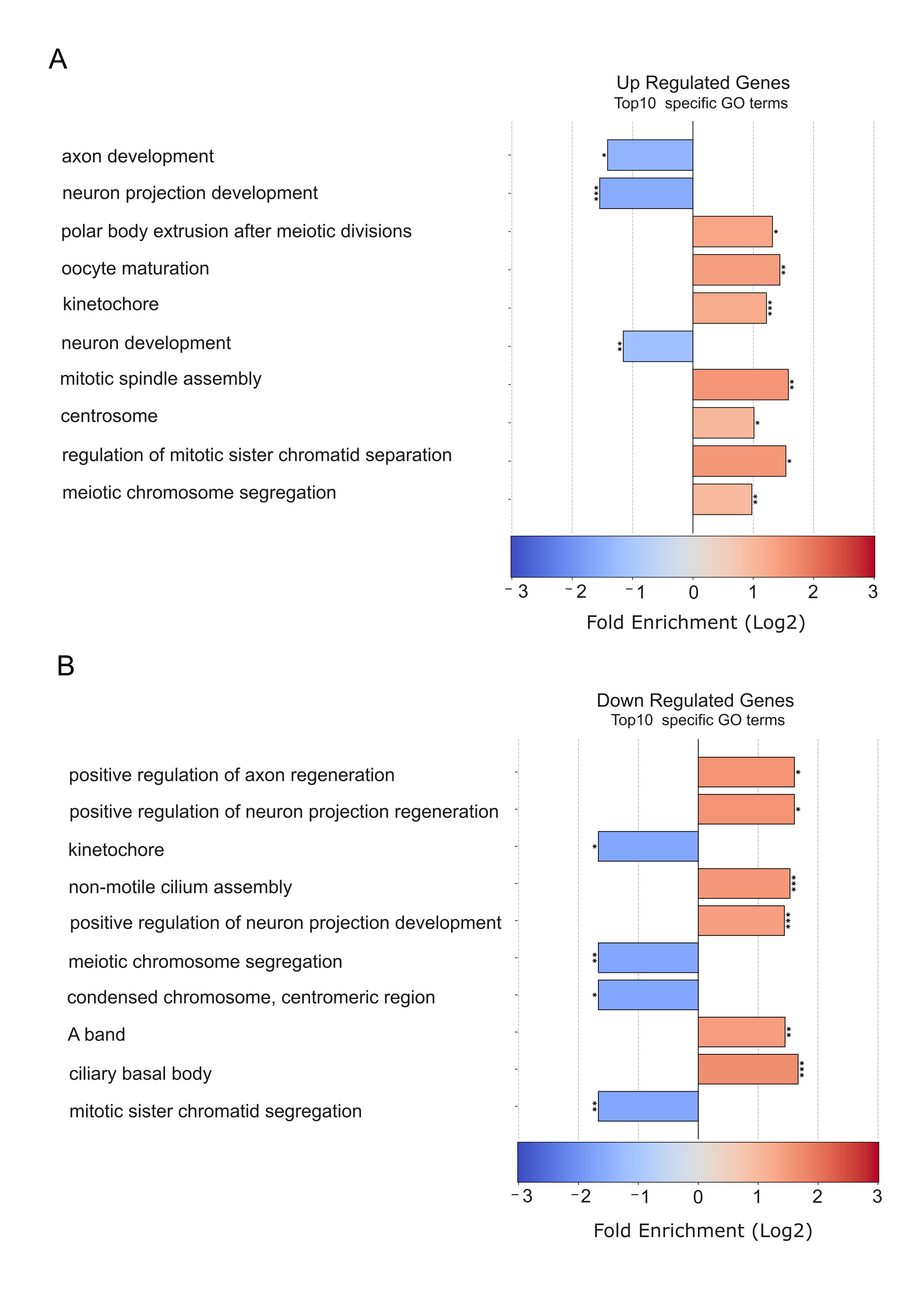

### Supplementary Figure3

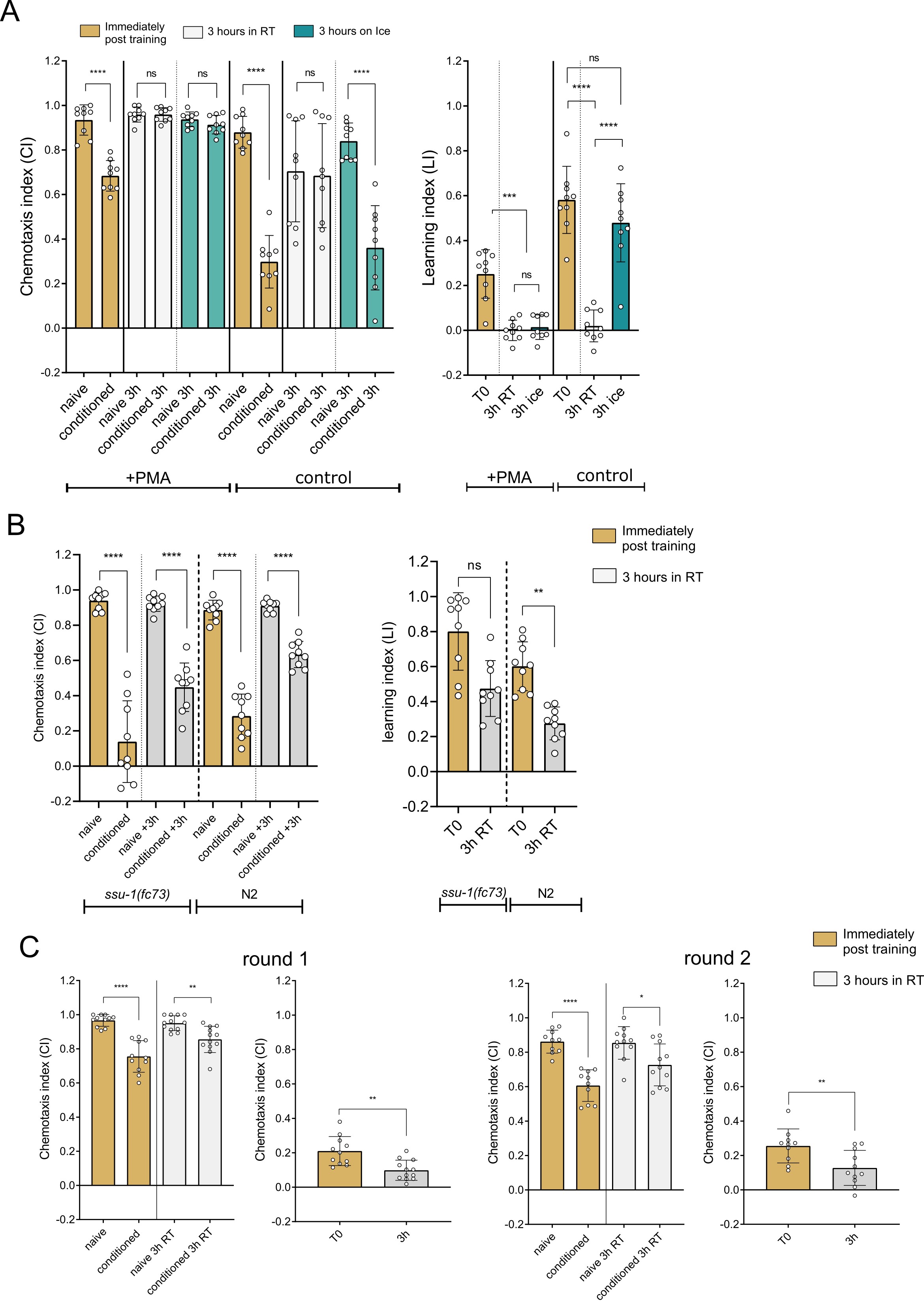

### Supplementary Figure4

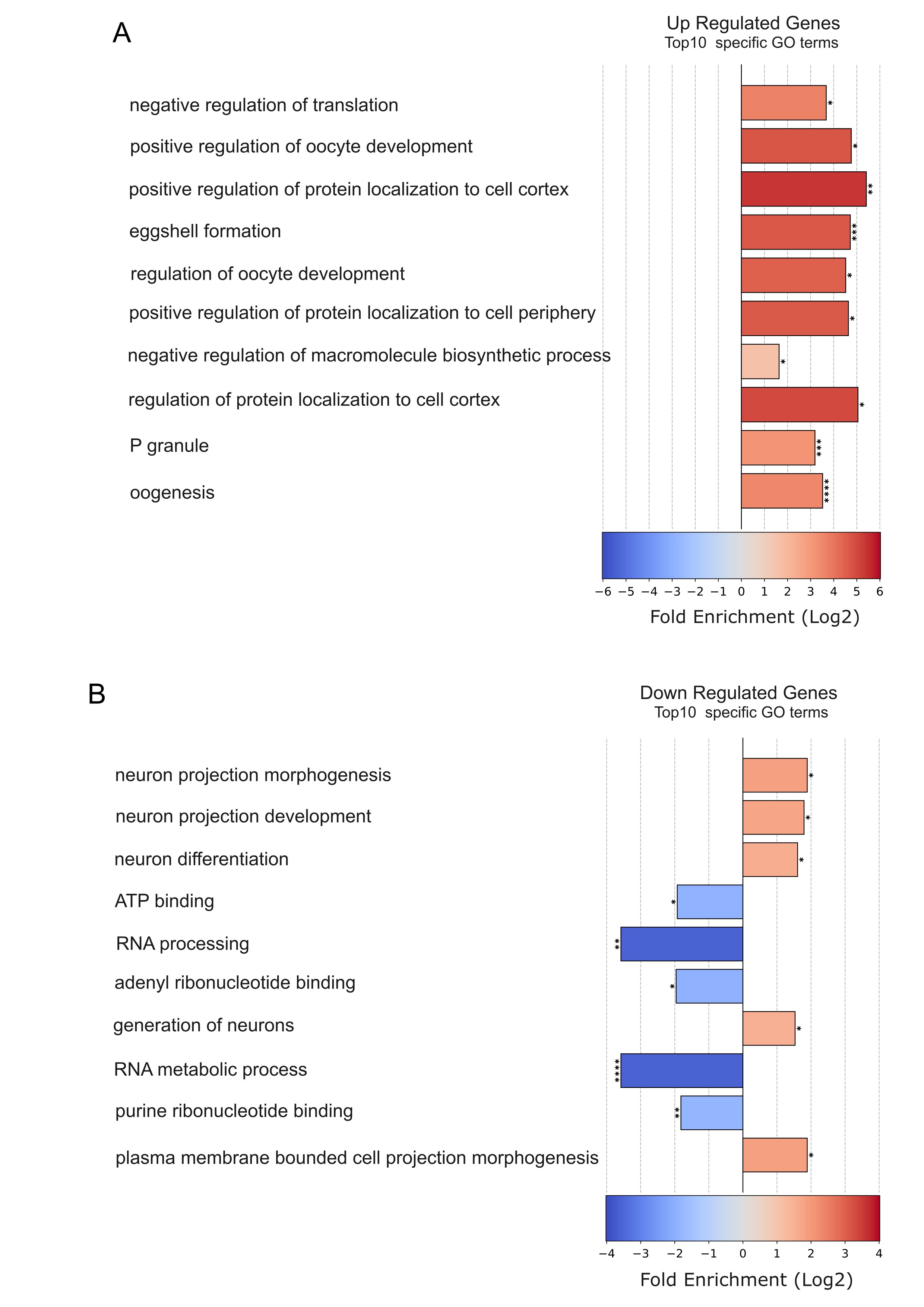

### Supplementary Figure5

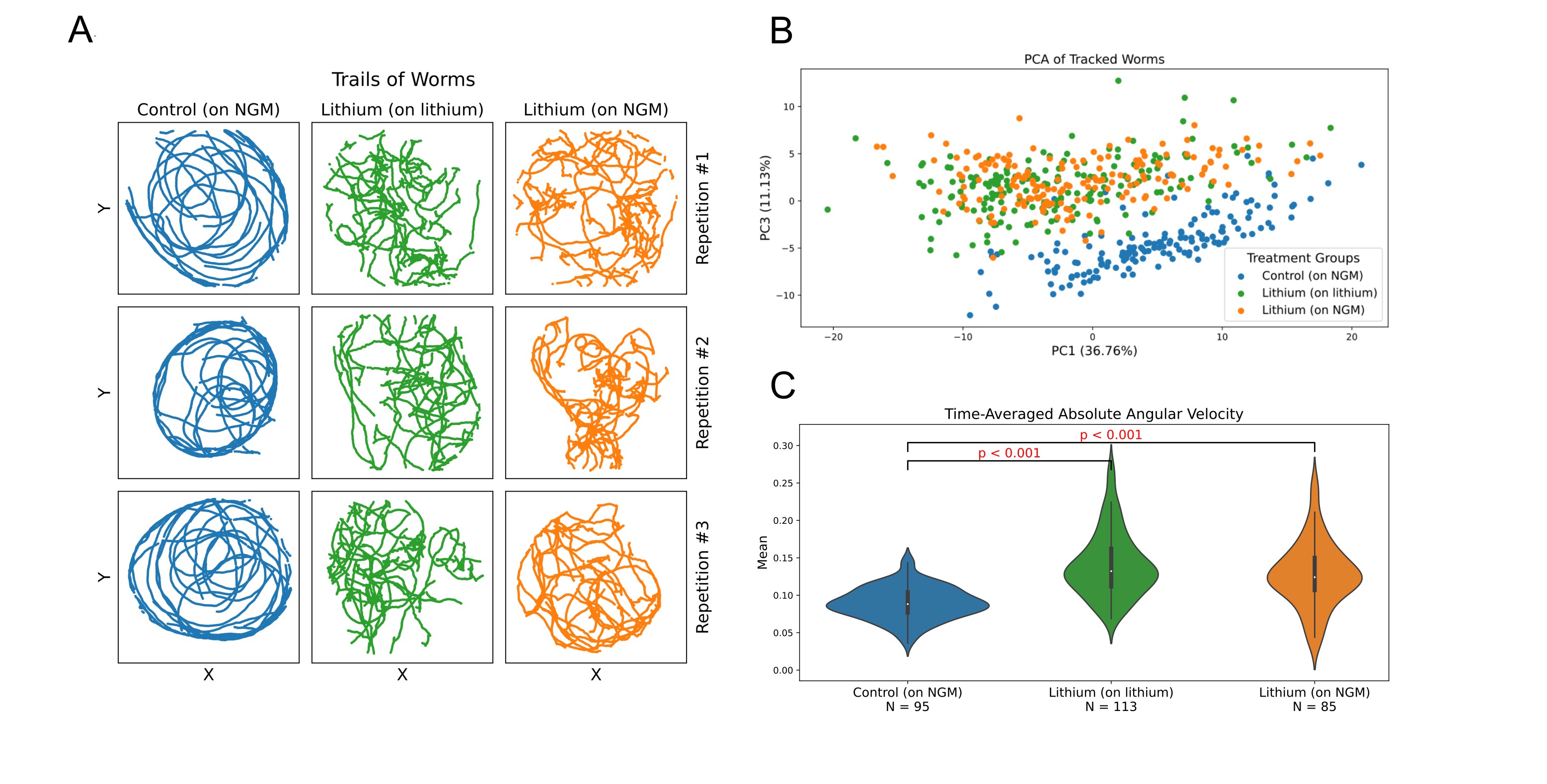
