## Supplementary Figure Legends for "A Tunable and Druggable Mechanism to Delay Forgetting of Olfactory Memories in *C. elegans*"

**Supplementary Figure 1. Forgetting of AWC sensed odor associations is delayed on ice.**

**A-B**: Chemotaxis outcomes for worms trained to different AWC sensed odors or control worms, assessed immediately after training or after a period of rest either on ice or at room temperature (RT). Left panel – Chemotaxis Index (CI); Right panel – Learning Index (LI). **A.** Worms trained for the odor 2,4-pentanedione (Pd). Statistical significance was calculated using a two-tailed Mann-Whitney test (CI), or ANOVA followed by Tukey's multiple comparisons test (LI) (N=12). **B.** Worms trained for the odor benzaldehyde, CI analyzed using a two-tailed Mann-Whitney test and LI analyzed using Kruskal Wallis followed by Dunn's multiple comparisons test (N from left to right, CI: 9,12,8,9,9,12; LI: 11,9,12. **C.** Worms trained for the odor butanone tested after 24h on ice. Statistical significance of the CI was calculated using a two-tailed Mann-Whitney test N=7. In all panels, The graphs comprised of 3 biological repeats with a minimum of 2 technical repeats in each. Each dot in the LI graph represents the difference between one conditioned plate and one naïve plate in the same biological repeat. Bar graphs denote mean ± standard deviation. Statistical significance is denoted as ns (not significant, $p > 0.05$), * ($p < 0.05$), ** ($p < 0.01$), *** ($p < 0.001$), **** ($p < 0.0001$)

**Supplementary Figure 2. Enrichment analysis of upregulated genes in worms that delay forgetting on ice show enrichment for germline functions and depletion of in neuronal functions.**

Displayed are the top 10 most specific (determined by their depth in the GO directed acyclic graph) experimentally derived, Gene Ontology (GO) annotations for genes that are either **A.** up-regulated or **B.** down-regulated in cold-sensitive worms while placed on ice compared to cold tolerant worms on ice. The presented list is comprehensive and has not undergone curation of any terms. Analysis and figure were generated using RNAlysis (Teichman et al. 2023) software tool.

**Supplementary Figure 3. Lithium delays forgetting independently of its described effect on the ASJ neurons.**

**A.** Chemotaxis outcome of worms treated with 0.05mg/ml of the DAG analog, PMA (Phorbol-12-myristate-13-acetate) for the 90min training session or control worms treated with ethanol as control during the same time frame. Worms were assessed immediately after training or after a period either on ice or at room temperature (RT). CI: two-tailed Mann-Whitney test. LI: one-way ANOVA followed by Tukey's multiple comparisons test N=9. **B.** Chemotaxis outcomes for trained and control *ssu-1*(fc73) mutants, or control WT (N2) treated overnight with 15mM of LiCl assessed immediately after training or after period either on ice or at room temperature (RT) Chemotaxis Index (CI), analyzed using a two-tailed Mann-Whitney test. Learning Index (LI): evaluated via Kruskal-Wallis test followed by Dunn's multiple comparisons test N=9. **C.** Chemotaxis outcomes for trained and control worms in worms expressing a caspase under the ASJ-specific *trx-1* promotor (VZ823 *jxEx100[pQZ37(Ptrx-1::ice) + Pofm-1::gfp)*]) assessed immediately after training or after period at room temperature (RT). This experiment was conducted twice to ensure reproducibility, denoted – round 1 and round 2 (each experiment was conducted with three biological repeats). Chemotaxis Index (CI), analyzed using a two-tailed unpaired t-test (round1 N from left to right CI: 11,11,12,12 LI: 11,12 round 2 N from left to right CI: 10,10,11,11 LI: 10,11). Learning Index (LI): two-tailed unpaired t-test. The graphs are comprised of 3 biological repeats with minimum of 3 technical repeat in each. Each dot in the LI graph represents the difference between one conditioned plate and one naïve plate in the same biological repeat. Bar graphs denote mean ± standard deviation. Statistical significance is denoted as ns (not significant, $p > 0.05$), * ($p < 0.05$), ** ($p < 0.01$), *** ($p < 0.001$), **** ($p < 0.0001$)

**Supplementary Figure 4. Enrichment analysis of genes that show a significant change in expression upon lithium treatment reveals patterns similar to those observed in worms that delay forgetting on ice.** Displayed are the top 10 most specific (determined by their depth in the GO directed acyclic graph), experimentally derived GO annotations for genes that are either **A.** up-regulated or **B.** down-regulated upon lithium treatment; We included only the genes that change specifically in cold sensitive worms but not in cold tolerant worms (as these are the genes that are likely to contribute to the ability of the worms to delay forgetting). The presented list is comprehensive and has not undergone curation of any terms. Analysis and figure were generated using RNAlysis software tool (Teichman et al. 2023).

**Supplementary Figure 5. Automated tracking of lithium-treated worms.**

**A.** Worms path reconstruction in 2D. Control worms (blue) were kept on regular NGM plates. One experimental group (green) was cultivated overnight on 15mM lithium plates and then tracked on 15mM lithium plates, the other experimental group (orange) was cultivated on 15mM lithium plates and tracked on control NGM plates (off lithium). **B.** PCA of worm trajectories. Each dot in the PCA space represents a single worm trajectory (time-averaged in absolute value). The percentage of variance represented by the first and third principal components, out of the total variance in the original high-dimensional space, is shown on the x-axis for PC1 and on the y-axis for PC3. **C.** Angular velocity changes in lithium-treated and control worms. Each dot represents a time-averaged absolute value of angular velocity of a single worm trajectory.
