## Supplementary Methods for "A Tunable and Druggable Mechanism to Delay Forgetting of Olfactory Memories in *C. elegans*"

### Strains:

ZD1687, mgIs40; *ssu-1(fc73)* gifted by Prof. Dennis Kim, ASJ ablation strain VZ823 *jxEx100[pQZ37(Ptrx-1::ice) + Pofm-1::gfp)*] gifted by Prof. Joy Alcedo.

### Olfactory training for benzaldehyde, and 2,4-pentanedione:

The protocol for olfactory training with benzaldehyde, and 2,4-pentanedione was adapted from Nuttley et al., 2002. Day-one adult worms were subjected to three washes in S-basal buffer before being transferred onto 90mm chemotaxis agar plates (detailed protocol available in the chemotaxis assay section). For the olfactory training, a 2 µl droplet of each undiluted odorant (or none for naïve group) was carefully placed onto a small square of parafilm which was positioned on the interior side of the assay plate lid, ensuring controlled exposure to the volatile compounds.

### GO enrichment analysis:

GO enrichment analysis was conducted using the RNAlysis software (Teichman, G. et al. 2023). The graph displays the 10 most specific GO terms (determined by their depth in the GO directed acyclic graph) that were found to be statistically significant. Enrichment analysis was performed using the classic annotation propagation algorithm, after filtering the GO annotations for experimentally-derived annotations only (evidence code: Inferred from Experiment, EXP). The background set for enrichment was defined as genes with detectable expression in at least one sample, with at least one GO annotation assigned to them. Upregulated and downregulated gene sets were determined based on differential expression analysis comparing cold-sensitive worms subjected to ice exposure with cold-tolerant worms subjected to ice.

To analyse changes in gene expression resulting from lithium treatment, we performed differential expression analysis, comparing worms treated overnight with 15mM LiCl to control worms maintained on NGM. We focused on genes that were differentially expressed in cold-sensitive worms, excluding those that changed in cold-tolerant worms. We identified genes whose expression was altered by lithium treatment specifically in the cold-sensitive worms that delay forgetting, but not in those that are cold-tolerant and do not delay forgetting.

**Worm tracking and analysis of behavioural changes:**

For analysis of worm's behaviour upon exposure to lithium, a small population of synced young adult worms were transferred to either 15mM LiCl or control NGM plate and kept there overnight. Then, 10 individual worms of the same treatment were transferred to a single NGM (either containing 15mM LiCl or control) plate seeded with fresh bacterial lawn and each plate was recorded for 10 minutes with 2fps (Niko worm tracker, Camera stage with adjustment mechanism, Sample stage for 60mm dish Backlighting Stage). Videos then underwent image analysis for feature extraction by the Tierpsy software (Javer et al., 2018), further manipulation and downstream computational analysis was performed using python scripts utilizing the output files made by Tierpsy analysis.

**Path Reconstruction**: The spatial trajectories of the worms were visualized using data obtained from the tracking software. This involved the utilization of the 'coord_x_body' and 'coord_y_body' features from the Tierpsy analysis.

**PCA Analysis** Samples with less than 80 frames were excluded to avoid noise stemming from short samples, subsequently we computed the time-averaged absolute values for each feature to preserve the integrity of directional data and applied standard scaling (Z-score normalization).

**Feature Comparison** similarly to the PCA analysis, trajectories less than 80 frames long were omitted. For the remaining data, we calculated the time-averaged absolute values of the angular velocity, statistical significance was calculated utilizing two-tailed Mann-Whitney U test.
